# The T cell receptor repertoire reflects the dynamics of the immune response to vaccination

**DOI:** 10.1101/2021.12.09.471735

**Authors:** Kevin Mohammed, Austin Meadows, Saboor Hekmaty, Sandra Hatem, Viviana Simon, Anitha D. Jayaprakash, Ravi Sachidanandam

## Abstract

Early, high-resolution metrics are needed to ascertain the immune response to vaccinations. The T cell receptor (TCR), a heterodimer of one α and one β chain, is a promising target, with the complete TCR repertoire reflecting the T cells present in an individual. To this end, we developed Tseek, an unbiased and accurate method for profiling the TCR repertoire by sequencing the TCR α and β chains and developing a suite of tools for repertoire analysis. An added advantage is the ability to non-invasively analyze T cells in peripheral blood mononuclear cells (PBMCs). Tseek and the analytical suite were used to explore the T cell response to both the COVID-19 mRNA vaccine (n=9) and the seasonal inactivated Influenza vaccine (n=5) at several time points. Neutralizing antibody titers were also measured in the covid vaccine samples. The COVID-19 vaccine elicited a broad T cell response involving multiple expanded clones, whereas the Influenza vaccine elicited a narrower response involving fewer clones. Many distinct T cell clones responded at each time point, over a month, providing temporal details lacking in the antibody measurements, especially before the antibodies are detectable. In individuals recovered from a SARS-CoV-2 infection, the first vaccine dose elicited a robust T cell response, while the second dose elicited a comparatively weaker response, indicating a saturation of the response. The physical symptoms experienced by the recipients immediately following the vaccinations were not indicative of the TCR/antibody responses. The TCR responses broadly presaged the antibody responses. We also found that the TCR repertoire acts as an individual fingerprint: donors of blood samples taken years apart could be identified solely based upon their TCR repertoire, hinting at other surprising uses the TCR repertoire may have. These results demonstrate the promise of TCR repertoire sequencing as an early and sensitive measure of the adaptive immune response to vaccination, which can help improve immunogen selection and optimize vaccine dosage and spacing between doses.

## Introduction

Vaccines provide prophylactic immunization against target viruses by inducing a lingering adaptive immune response that creates immunological memory^1^, which protects against future infections. The adaptive immune response consists of the T cell-mediated response and the humoral antibody-response (mediated by B cells). In the antibody response, B cells activated by the vaccine immunogen differentiate into plasma cells capable of producing neutralizing antibodies, which can bind to proteins on the target virus and hinder its infectiousness/virulence, or memory B cells primed to become plasma cells upon reinfection of the target virus. The cell-mediated response consists of activated effector CD4 T cells, which induce B cells to produce antibodies or recruit the microbicidal functions of other immune cells including macrophages; activated effector CD8 T cells which destroy virus-infected cells; and memory T cells (CD4 or CD8), which persist long after vaccination and are primed to become effector T cells. Other immune cells can exhibit memory-like behavior as well, such as NK cells^2^.

Vaccines come in many forms, as inactivated viruses, viral protein fragments, vectors that infect cells to produce viral proteins, or mRNA that is transcribed in cells to produce viral proteins. Vaccines aim to introduce pathogenic protein subunits to trigger a robust B and T cell (cellular) response^3^, involving B and T cells targeting multiple epitopes, which results in a robust, broad antibody (humoral) response. The T cells reflect an early response that sets stage for the later B cell/antibody response^4^. The reported efficacies for these vaccines can range from 30% for certain flu vaccines, to 90% for the Covid mRNA vaccines^5,6^. The vaccine against Influenza, a rapidly-evolving, negative-single-strand, segmented RNA virus, is comprised of inactivated viruses, in contrast to the mRNA vaccines, which induce intracellular production of viral proteins or their sub-units. The mRNA vaccines likely induce both CD4 and CD8 T cells^7,8^, since the viral proteins are produced intracellularly, while the flu vaccines most likely induce only CD4 T cells^9^. It is not a given that mRNA vaccines work, an early attempt to use mRNA vaccines for Rabies failed^10^.

The covid-19 pandemic, caused by the SARS-CoV-2 virus, has set off a scramble to develop vaccines, validate their efficacy and administer them around the world. This has brought to the forefront the importance of measuring the response to vaccines and the methods of doing so^11,12^. Epidemiological data is often the final arbiter of responsiveness to vaccines, but the time and effort required to collect and collate the data does not allow nimble modification and testing of vaccination strategies. The response to vaccines can also be studied in individuals, by detecting antibodies against viral proteins from serum, most frequently by using an Elisa test^13,14^ or by detecting activated B and/or T cells.

One drawback of the antibody-based assays is that it takes a few weeks after vaccination to develop detectable levels of antibodies^15^. These assays might not detect low levels of antibodies that might be sufficient to provide protection, especially long after the infection or vaccination. The assays only detect antibodies against particular viral proteins/peptides, missing antibodies against other parts of the viral proteome which might provide adequate protection^16^. An alternative to antibody detection is to identify T cells and B cells that are responsive to the vaccine, using assays such as the ELISpot, to measure release of either interferon gamma or granzyme B from the activated cells^17^. ELISpot indirectly correlates activity to specificity, is not always accurate and can occasionally mislead when the cells aren’t activated despite possessing the antigen-specific receptors^18^.

We hypothesized that the TCR repertoire could add an important new dimension to the measurement of the response to vaccines, as it would identify the T cell receptors (TCR) of clones that respond to the vaccination^19,20^. The ability to use non-invasive methods, such as sampling the PBMCs, would be advantageous as well. Each T cell has on its surface several copies of the TCR that uniquely identifies it, which bind to specific viral peptides derived from the vaccine immunogen presented by Human Leukocyte Antigen (HLA) molecules on the surface of antigen presenting cells. The peptide chosen for presentation is determined by the HLA, while the binding efficiency is determined by the TCR and the peptide. Successful binding induces clonal expansion of the T cell.

The TCR is a heterodimer of two trans-membrane polypeptide chains (TCRα and TCRβ) linked by covalent disulfide bonds. Germline TCRα and TCRβ loci undergo rearrangement during the development of each T cell, wherein one of several V, D (only for TCRβ), and J regions are joined, accompanied by mutations at the joints^21^, which define the complementarity determining region CDR3. There are two additional hyper-variable complementarity determining regions (CDR1, CDR2) located on the TCRα and TCRβ chains, which together determine the binding specificity of the TCR to peptide-HLA complexes, but the CDR3 contributes the most to antigen specificity^22^. Thus, we reasoned that sequencing the TCR present in an individual should yield a profile of the relative abundances of vaccine-induced T cell clones, identified by their V, J and CDR3 sequences, offering a more detailed view into the adaptive response to vaccinations in addition to the traditional, antibody-centered approaches. To this end, we developed Tseek^23^ for unbiased, sensitive profiling of TCRα and TCRβ chains (**Fig. 1, Methods**).

**Figure 1.**
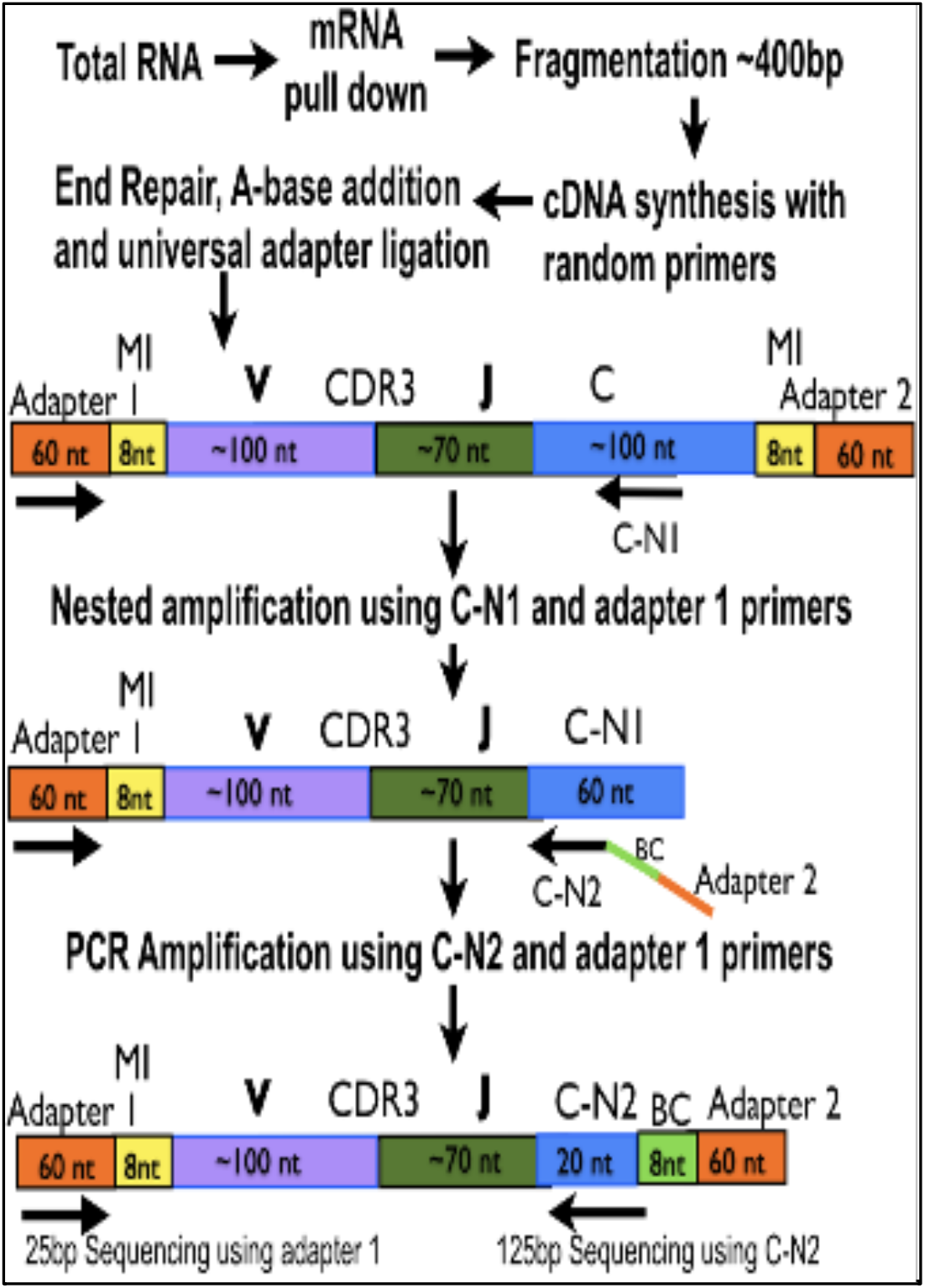
Tseek protocol. RNA-based approach that does not rely on prior knowledge of the V-segments. The key steps are 1) the fragmentation of mRNA, and ligation of adapters to cDNA synthesized from the fragments, 2) amplification with a primer from the C-region on the 3’ end (C-N1, one each for α, β) and the universal 5’ adapter (adapter 1), 3) A nested PCR using adapter 1 on the 5’ end and a second 3’ primer in the C (C-N2) to which is attached barcodes and Adapter 2, resulting in the final sequence. This is then sequenced using the C-N2 primer.

**Figure 2.**
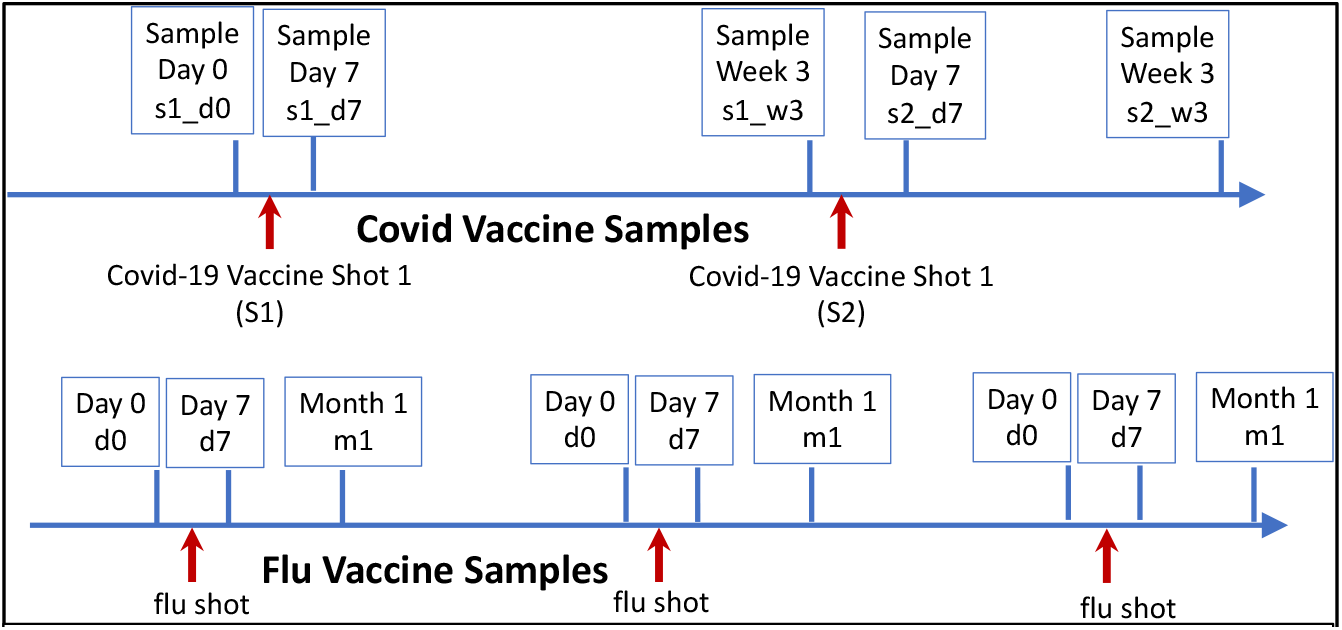
Vaccination and sample collection schedules. Blood samples were collected from individuals who received the Covid-19 mRNA vaccine (top) and flu vaccines (bottom) according to the schedules shown in the figure. For the two Covid doses, a day 7 (d7) and week 3 (w3) response after each dose were measured. For the flu doses, given once per year over several flu seasons, multiple blood draws, before the dose, at day 7 (d7), and at one-month (m1) after the dose, as well as a 6-month measurement. The s1_d0 samples were missed in several covid vaccine samples

To explore the utility of Tseek in evaluating vaccine responses, we compared the responses to mRNA COVID-19 vaccines and the annual influenza vaccines. The two vaccines occupy different landscapes; the efficacy of the COVID-19 vaccine is high (∼ 90%) while that of the influenza vaccine is low (reaching 30% in some seasons), based on epidemiological and antibody data^5,6^ and the modality of delivery is also different (inactivated virus versus mRNA). We wanted to establish the Tseek data from PBMCs reflected the different outcomes of the vaccines. To do so, we utilized PBMC samples from people inoculated with the yearly influenza vaccine over several flu seasons, spanning up to several years. We also collected PBMC samples from people who were inoculated twice with the mRNA vaccine at 5 time points throughout the vaccination course (**Table 1**). For the individuals in the COVID-19 group, we also measured the levels of neutralizing antibodies against SARS-CoV-2 spike protein, to correlate with the T cell response.

**TABLE 1.** Covid Vaccine cohort table. Qualitative measures of the TCR and antibody response to the covid mRNA vaccine (derived from data in **Table 2**) as well as the symptoms experienced by the donors after their vaccination doses are shown. The high, medium and low categories were defined by ranking the values and roughly grouping the results into natural clusters, this could change as more data is accumulated. B3 and B4 have recovered from a previous covid infection. There is no correlation between the symptoms immediately after the vaccine dose and the adaptive immune response. Higher TCR responses did not track with the antibody response, likely because only a subset of the TCR response is responsible for the antibody response. But a low TCR response (B1, B6, B8) seems predictive of a low antibody response, and the one case (B9) with very low TCR response also has little antibody response. B5, B8 and B9 had the d0 samples, allowing us to infer patterns for the reactive clones in the others which did not have the d0 samples. *B8 received the Moderna mRNA vaccine, the rest received the Pfizer vaccine.

| Donor | Gender | Prior Covid | Antibody response | TCR response | Symptoms Dose1 | Symptoms Dose 2 |
| --- | --- | --- | --- | --- | --- | --- |
| B1 | Male | No | low | low | Mild | Mild |
| B2 | Female | No | medium | high | None | None |
| B3 | Male | Yes | high | medium | None | None |
| B4 | Female | Yes | medium | medium | None | None |
| B5 | Male | No | high | medium | Mild | Severe |
| B6 | Male | No | low | low | Mild | Mild |
| B7 | Male | No | medium | high | None | None |
| B8* | Male | No | low | low | None | Severe |
| B9 | Female | No | very low | very low | None | None |

This study establishes the feasibility and utility of using PBMCs to monitor the changes in the TCR repertoire in response to vaccines.

## Results

### Tseek is an accurate, unbiased method of profiling the complete TCR repertoire

We developed Tseek, for unbiased, sensitive profiling of the α and β TCR chains in bulk (**Fig. 1**). It uses RNA, instead of DNA and does not require prior knowledge of the V-segments, needed by most other methods, allowing its use in non-model animals such as swine^24^. Tseek can work with total RNA from any type of material, it does not need T cells to be isolated and is sensitive to small numbers of T cells in the sample.

Many alternative methods of sequencing the complete TCR repertoire rely on several dozen polymerase chain reactions (PCRs) primed off selected V segments (70 for TCRβ and 52 for TCRβ in humans^25^), which result in biases due to the use of myriad V-primers in complex multiplex PCR reactions. Tseek (**Fig. 1**) provides an unbiased approach, which has been used for profiling TCR repertoires from mammals including human, mice, swine, dogs, cats, and monkeys) and its use has been demonstrated in several published studies^24,26,27^. The only other comparable unbiased method uses RACE-PCR (kit from Takara). In a head-to-head comparison, based on preparing the same sample (from another study^28^ kindly provided by the authors) by both methods, Tseek showed good concordance with RACE-PCR, while offering higher sensitivity and specificity(**Fig. S1A**). We also observed good concordance between Tseek and qPCR for the expression levels of selected V segments (**Fig. S1B**).

### The TCR repertoire is a fingerprint, specific to an individual

We used the Jensen-Shannon (JS) metric (Methods section) to define a distance between samples based on our TCR CDR3 measurements. Dendrograms were constructed using hierarchical clustering based on the pairwise JS distances, without relying on sample identifiers. In the Covid-19 (**Fig. 3A, S3A)** and the Influenza (**Fig. 3B, S3B**) vaccine sets, the samples clustered by donor for the **α** and **β** CDR3 despite the samples being taken years apart in some cases. This suggests that 1) the adaptive response to vaccinations (and infections) are a perturbation on the overall TCR profile, and 2) there are a set of unperturbed clones characteristic for each individual, providing a fingerprint for identification of the donor based solely on this repertoire. This provides a metric to distinguish identical twins.

**Figure 3.**
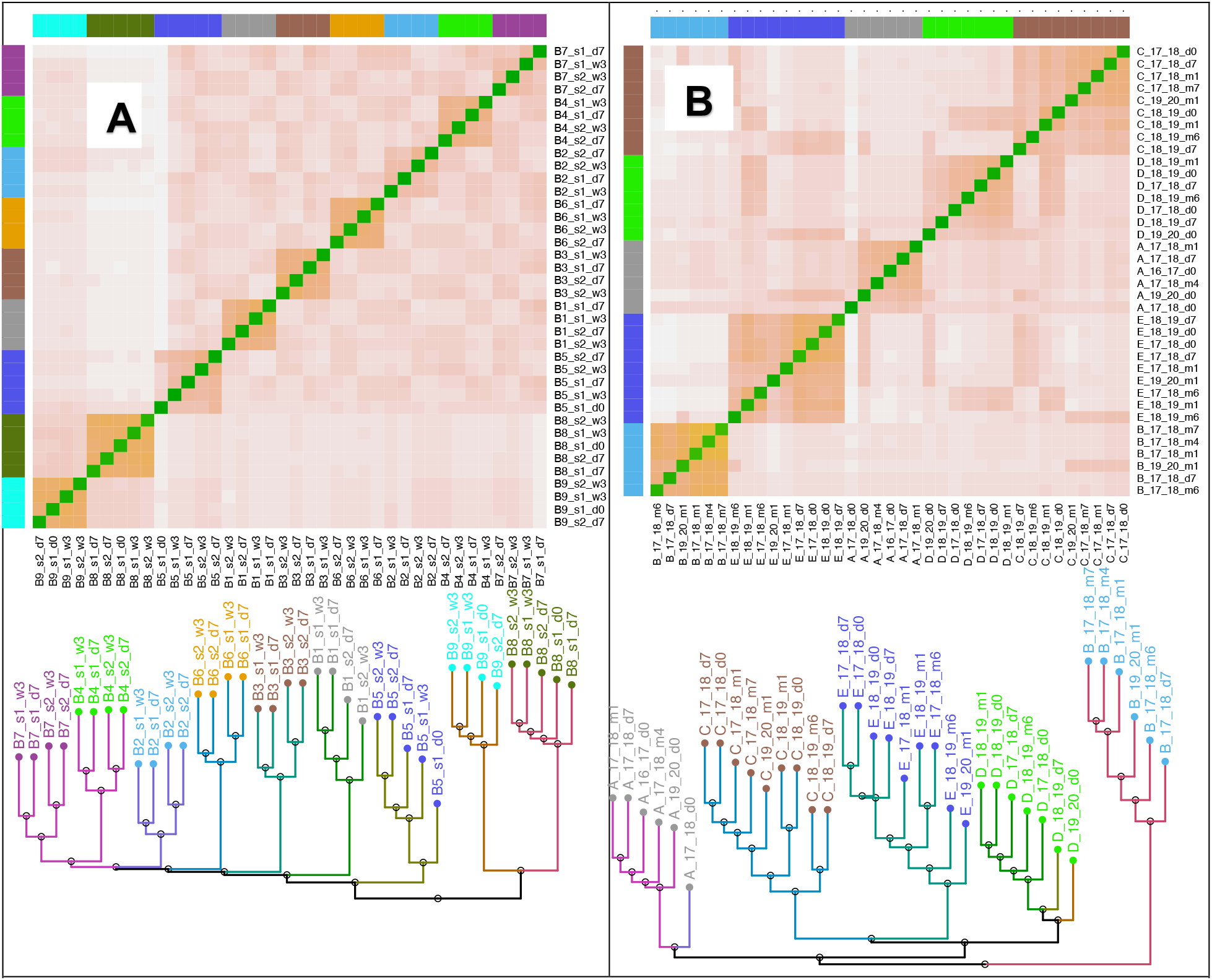
Clustering of T cell repertoire (CDR3) from the Covid/Flu vaccine samples. Distances between samples were defined for the CDR3 data using the Jensen-Shannon metric (an information-theoretic measure, Supplementary material). The heatmaps show the distance between pairs of samples, while the dendrograms show the samples cluster by individual. **A) Covid CDR3 β**. The first dose is s1, the second dose is s2, d0 is the pre-vaccine sample, d7 is 7 days post vaccination, w3 is 3 weeks post vaccination. For each individual, the two samples after the second vaccine dose (s2_d7 and s2_w3), cluster separately from the two samples after the first vaccine dose (s1_d7 and s1_w3) suggesting distinct sets of T cells react to the two doses. The second dose broadens viral epitopes targeted, likely improving the immunity provided by the vaccination. (**Fig. 3A** shows the corresponding α tree) **B) Flu β CDR3** 19_20 refers to the 2019-2020 flu season, d0 is the pre-vaccine sample, d7 is 7 days post vaccination, m1 is 1 month post vaccination. The clustering by vaccine dose does not occur in some individuals (B, D), and the m4 and m6 samples seems to have lost “memory” of the dose, the T cell response does not persist beyond 2-3 months. Even though there are responsive clones for each flu season (Table 2A, B), the relative lack of clustering of the tree by vaccine dose, in contrast to the covid vaccine data, suggests that the response is “weaker” and has more individual and seasonal variability. **Fig. S3** shows data for the β.

### The TCR α and β chain repertoires are both responsive to vaccines

Because the TCR receptor is formed as a pair of an α and β chain, one would expect the changes in the two profiles should track each other. But we find that while the trees for **α** and **β** chains are similar, the numbers of responsive clones in the two chains are different, the responsive clones for **α** chain repertoire are more numerous compared to those for the **β** chain repertoire (**Fig. 4, 5**). This suggests the effect of the binding of the **α** and **β** chain in the dimer to the epitope are independent from one another and the effect of the binding of the two to the epitope is likely additive. Thus, both chains should be considered while evaluating the T cell response. Deciphering the implications of the differences in the response of the two chains needs a more controlled study, such as one using antigens in isogenic mice with defined MHCs.

**Figure 4.**
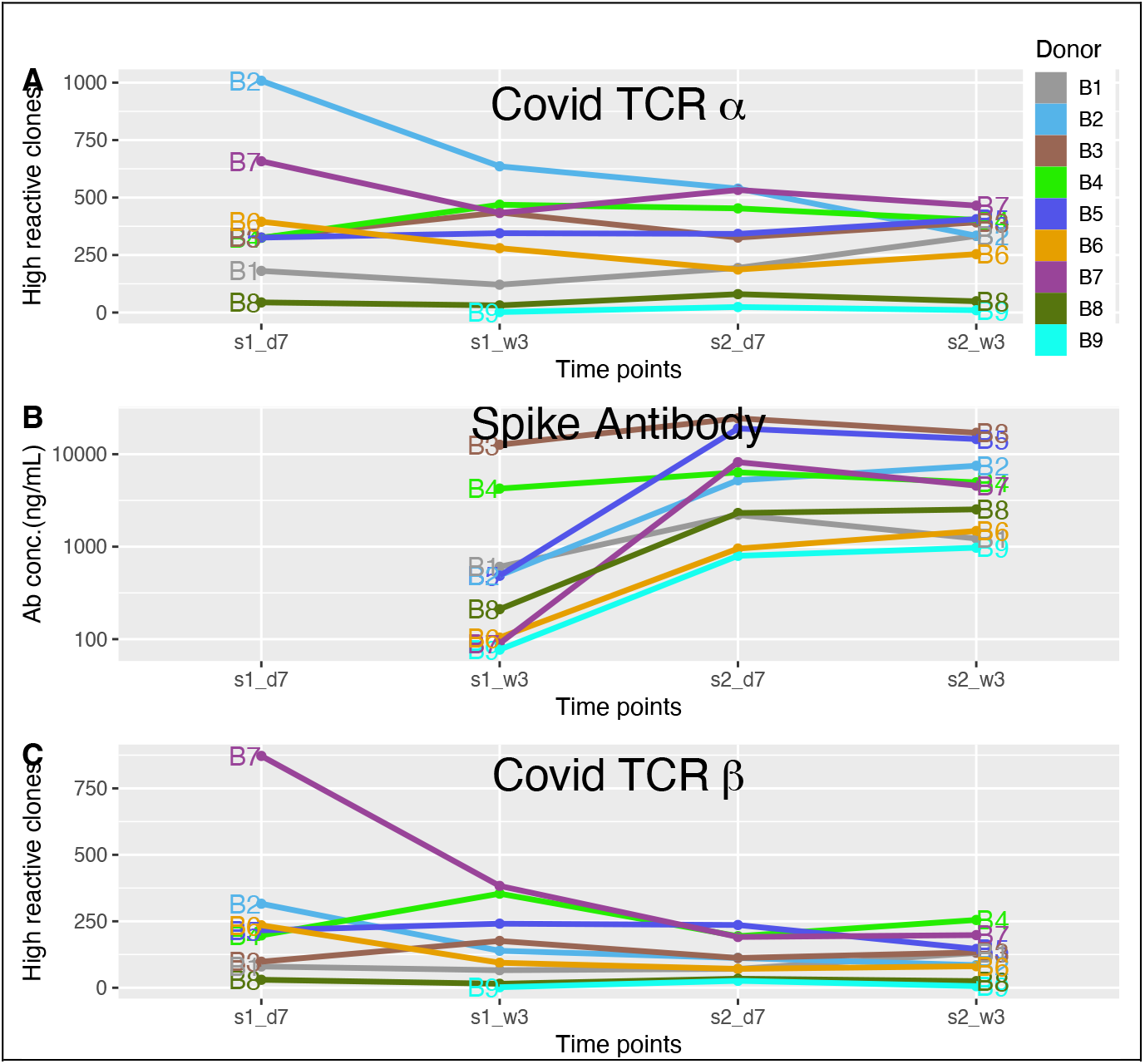
TCR and antibody response to the covid vaccine. There are nine individuals (donors) in this cohort who have received two covid mRNA vaccine doses (s1 and s2), approximately 3 weeks apart (day 7 or d7 and week 3 or w3). T cell receptors (bulk α and β) and the covid spike antibodies were measured at day 7 and week 3 post vaccination after each dose. At each time point, reactive clones (using r-fold_m_=10, ΔΔ 250, τ = 1500) that were H are counted, based on the H/L (high/low) classification. **A)** The plot shows counts of reactive α clones that are high (H) (y-axis) at each time point (x-axis). **B)** The antibody levels (y-axis, log-scale) at each time point (x-axis), and **C)** The high reactive β clones (y-axis) in each sample (x-axis). There are large individual variances in the response. B3 and B4 recovered from covid-19 infection months before the vaccine, both started out with high antibodies at s1_w3 that increased at s2_d7, but slightly dipped at s2_w3, while remaining high. The number of responsive clones is not as high as the others and is lower after s2, suggesting a saturation of the response, the infection seems to act like a first dose. The saturation of the T cell response in these individuals might reflect the fact that only a fixed number of epitopes from the spike protein could be presented by an individual’s MHC. B9 had a very low TCR response and almost no spike antibodies even after the second dose. B1, B6 and B8 have low TCR response compared to the others, and the antibody response after dose 2 also remains low.

**Figure 5.**
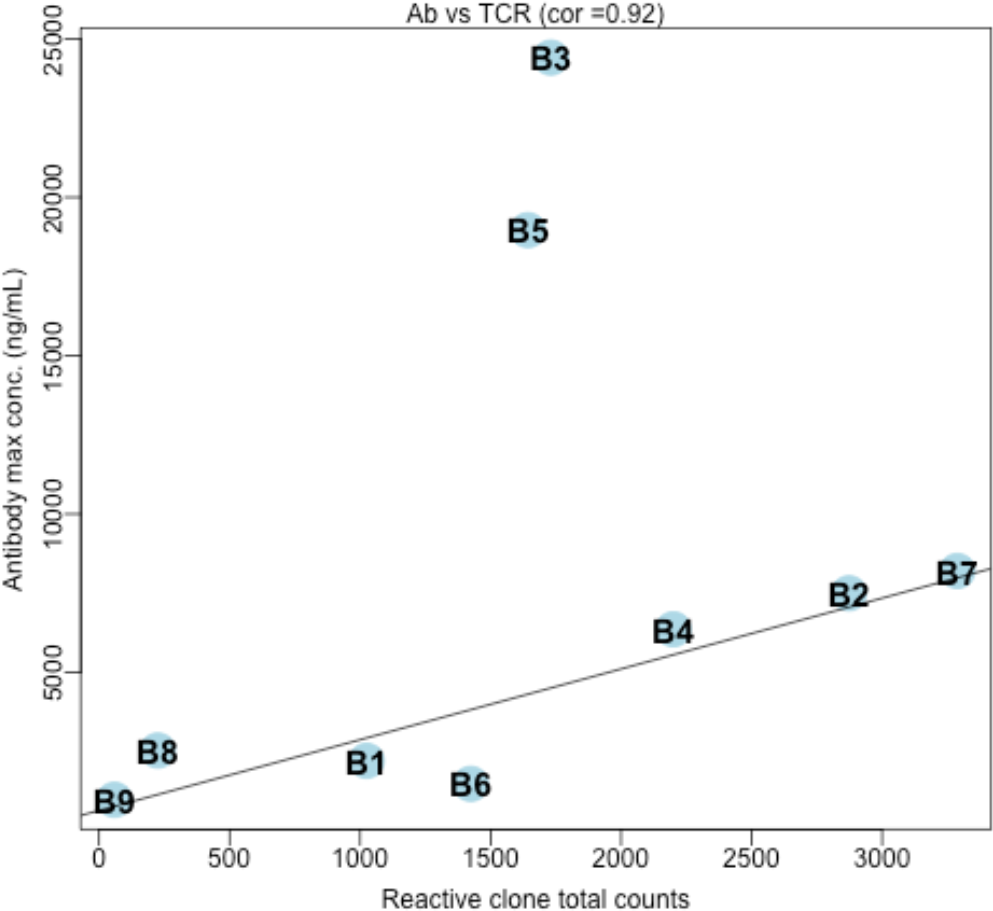
Antibody response (y axis – ng/mL) and reactive T cell clones (x axis – counts). The maximum of antibody levels is plotted against the total number of reactive clones measured in each donor, using data from **Table 2**. There is a strong correlation (0.92) between the T cell response and the subsequent antibody response. The linear fit and the correlation were calculated excluding the outliers, B3 and B5, who have a very strong antibody response.

### Landscape of the T cell response to the Covid-19 Vaccine

We collected two samples per vaccine dose, taken 1 week and 3 weeks after the dose (**Fig. 2**). Most clones were high only at one of the samples, falling below threshold in the other samples. This fact allowed us to identify responsive clones in donors missing the pre-vaccination sample. Despite the expanded clones being different at each sample, the repertoires, especially for the samples taken after dose 2, clustered together by the preceding vaccination dose, suggesting they share several expanded clones that are below the threshold in one or both samples. This is true for every individual in the study, despite individual differences in the response, suggesting that responsive

### Reactive clones and their classification

We define reactive clones as those that are clonally amplified in at least one of the samples (exact definitions are given in the Methods section). There are three parameters that determine if a clone is reactive, the fold change r-fold, which depends on Δ and needs to be above r-fold_m_ and the maximum frequency across samples for the clone, which needs to cross the threshold τ. We used, for the covid samples, r-fold_m_ = 10, Δ = 250 and τ = 1500 and for the flu samples, r-fold_m_ = 5, Δ = 300 and τ = 1000 A reactive clone can be either high (H) or low (L) at a timepoint (sample) depending on if its frequency is above or below the threshold (τ). The labels (**H** and **L)** form a binary code that can be stitched together into string codes for each clone. For ***n*** time samples there are 2^n^ possible string codes (e.g., **LLLH, HLLL** etc. resulting in 16 codes for the four samples). A clone with the string code “LHLL” is high only at time point 2. Grouping the reactive clones into classes based on their string codes yields the results shown in

**Table 2** and **Table 3**. For further insights, the journey of the clones over time can be represented by a directed acyclic graph (DAG, **Fig. 7**, Supplementary Material), since there are no cyclic (“going-back”) paths followed by the clones. We note several features in the data that are not sensitive to the exact values of r-fold_m_ and τ (**Fig.4)**. s1 and s2 stand for the two vaccination doses, and d7 stands for day 7 after the dose while w3 stands for week 3 after the dose.

a. The doses s1 and s2 induce distinct sets of high (H) reactive clones, with almost no shared clones between the sets (as seen in the DAG in **Fig. 7**)
b. Reactive clones high at day 7 (d7) are distinct from the ones high at week 3 (w3) for s1 and s2.
c. A direct consequence of **a)** and **b)** is that HLLL, LHLL, LLHL, LLLH are the dominant classes, few clones have more than one H in their string code.
d. B3 and B4, who had recovered from Covid-19, show reduced response after s2 relative to s1 (both LLLH and LLHL have fewer clones compared to clones classified as LHLL and HLLL). This suggests,
  i. There is a saturation of response to the second dose for these individuals
  ii. Covid-19 infection acts like a first dose of the vaccine in terms of the T cell response
  iii. A third booster dose for uninfected people would likely induce a weaker response compared to the original response.
e. Broadly, the T cell response (based on number of responsive clones) are indicative of the subsequent antibody responses (**Fig. 6, Table 1,2**). One individual (B9) with very poor T cell response seemed to have a minimal antibody response.

**Table 2.** TCR and antibody (Ab) response to the covid vaccine. There are nine individuals (donors) in this cohort who have received two covid mRNA vaccine doses (s1 and s2), approximately 3 weeks apart. T cell receptors (counts for bulk **α** and **β**) and the covid spike antibodies (ng/mL) were measured at day 7 and week 3 post vaccination after each dose. Reactive clones (using r-fold_m_=10, Δ =250,τ=1500) are represented by a string label, based on the H/L (high/low) classification at the four measurements (s1_d7, s1_w3, s2_d7, and s2_w3). Missing measurement (like the s1_d7 sample for B9), were represented with L. The TCR numbers show the αβ chain response. The last two columns are the highest spike antibody measurement and the number of reactive clones (sum of α and β for each individual. The responses can be sorted into high, medium and low for antibody levels (high > 10,000, 4000 < medium < 10,000, low < 4000), and for the TCR response (high > 2500, 2500 > medium > 1500, 1500 > low). **1)** Most reactive clones are high at only one sample point (so HLLL, LHLL, LLHL and LLLH dominate). **2)** B3 and B4 were infected with covid prior to vaccines and their spike antibody levels at s1_d7 reflect that, and their antibody levels seem to go down slightly after dose 2. **3)** B9 had almost no TCR response and almost no spike antibodies even after the second dose, and **4)** Mostly the TCR response presaged the antibody response (**Fig. 6**), the ones with very high levels of antibodies do not directly correlate with the number of reactive clones, likely because only a subset of reactive clones induce the antibody response. The graph in **Fig. 4** show the number of clones with H in each sample. * B9 is missing one measurement (s1_d7).

| Donor | Ab<br>s1_w3 | Ab<br>s2_d7 | Ab<br>s2_w3 | HLLL<br>$\alpha:\beta$ | LHLL<br>$\alpha:\beta$ | LLLH<br>$\alpha:\beta$ | LLHL<br>$\alpha:\beta$ | LLHH<br>$\alpha:\beta$ | HHLL<br>$\alpha:\beta$ | Ab<br>max | TCR<br>total |
| --- | --- | --- | --- | --- | --- | --- | --- | --- | --- | --- | --- |
| B1 | 606 | 2205 | 1226 | 130:62 | 70:46 | 288:111 | 141:58 | 19:5 | 18:6 | 2205 | 1025 |
| B2 | 488 | 5228 | 7497 | 892:289 | 506:108 | 232:62 | 423:90 | 34:12 | 53:19 | 7497 | 2873 |
| B3 | 12650 | 24412 | 17037 | 243:77 | 350:155 | 292:91 | 219:67 | 52:33 | 36:11 | 24412 | 1731 |
| B4 | 4246 | 6359 | 4962 | 207:124 | 340:273 | 282:161 | 314:115 | 51:38 | 50:28 | 6359 | 2200 |
| B5 | 485 | 18962 | 14577 | 188:130 | 200:150 | 238:63 | 180:143 | 37:16 | 19:18 | 18962 | 1643 |
| B6 | 104 | 954 | 1477 | 338:208 | 216:69 | 197:62 | 129:54 | 26:7 | 30:16 | 1477 | 1424 |
| B7 | 90 | 8199 | 4566 | 581:807 | 304:276 | 335:108 | 404:106 | 62:34 | 47:43 | 8199 | 3288 |
| B8 | 211 | 2312 | 2530 | 18:16 | 19:11 | 30:10 | 49:15 | 4:4 | 2:0 | 2530 | 225 |
| B9 | 77 | 796 | 975 | 0:0 | 0:1 | 6:2 | 19:21 | 3:4 | 0:0 | 975 | 59* |

**Table 3.**
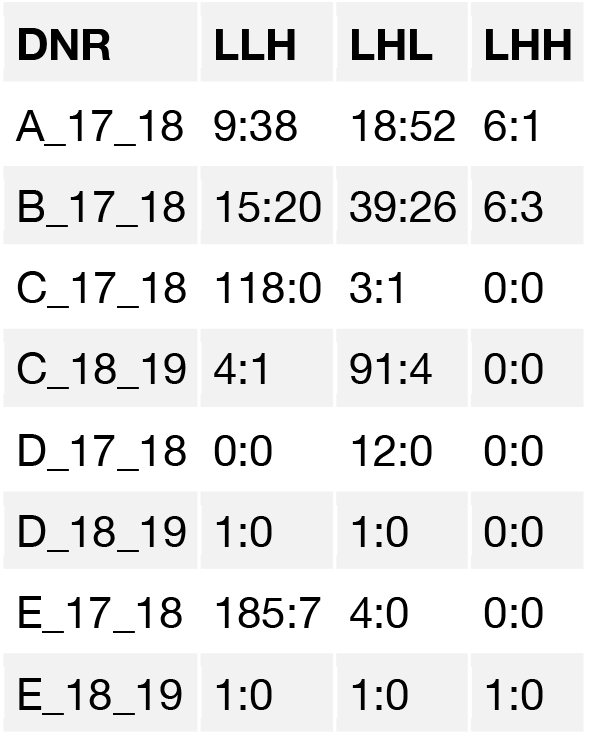
Flu Vaccine T cell response. There are five individuals in this cohort (A.. E), and they have received flu vaccine doses over several flu seasons (2017-2018, 2018-2019, and 2019-2020). In each case, the pre-vaccination (d0), the 1-week post vaccination (d7) and 1 month post vaccination (m1) measurements were used, measurements further out (months 4, 6, 7) have no signal. Reactive clones (using r-fold_m_=5, Δ =250, τ=1000) are represented by a string label, based on the H/L (high/low) classification at each measurement. Missing measurements (like the 1-week sample in B_19_20), are represented by L. The numbers show the αβ chain response. The response to flu vaccines is narrow compared to the response to covid vaccines, based on the clustering of the trees and the lower thresholds used to identify reactive clones. There is no coincidence between high reactive clones in week 1 versus month 1. Four individuals have a response (A, B, C and E) at least once and C has a response a year later too, suggesting immunologic memory is not the reason for non-responsiveness of some individuals. **Fig. 5** reflects the data in this table, showing counts of H at each time point.

**Figure 6.**
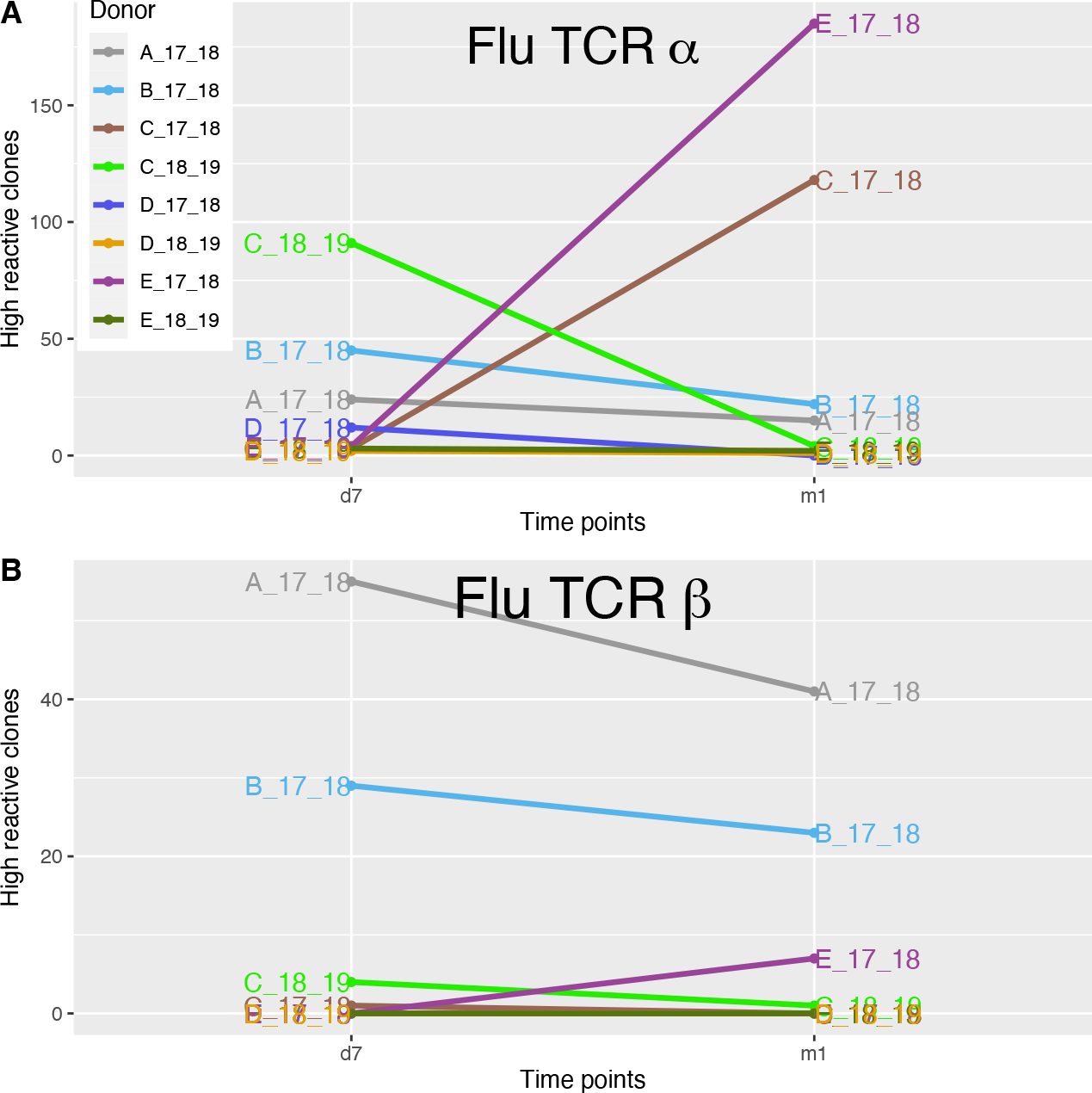
TCR response to the Flu vaccine. There are five individuals in this cohort (A, B, C, D, E), who have received flu doses over several flu seasons (2017-18, 2018-19, and 2019-20). In each case, the TCR response at pre-vaccination (d0), 1-week post vaccination (d7) and 1 month post vaccination (m1) were recorded. Reactive clones (using r-fold_m_=5, Δ=250 τ=1000) were identified and the number of clones that are expressed highly (H) at each time point are shown on the graph. **A)** The α chain cdr3 response. **B)** The β chain CDR3 response. The two responses seem decoupled. The overall response to flu vaccines is narrow compared to the response to covid vaccines, based on the clustering of the trees and the lower thresholds used to identify reactive clones. Four individuals have responses (A, B. C and E) and C has a response a year later too, suggesting immunologic memory is not the reason for non-responsiveness of some individuals. The T cell response could serve as a marker for who is benefiting from the vaccine and likely to be protected, but a larger study, correlating the data with antibody data is needed to develop a metric.

**Figure 7.**
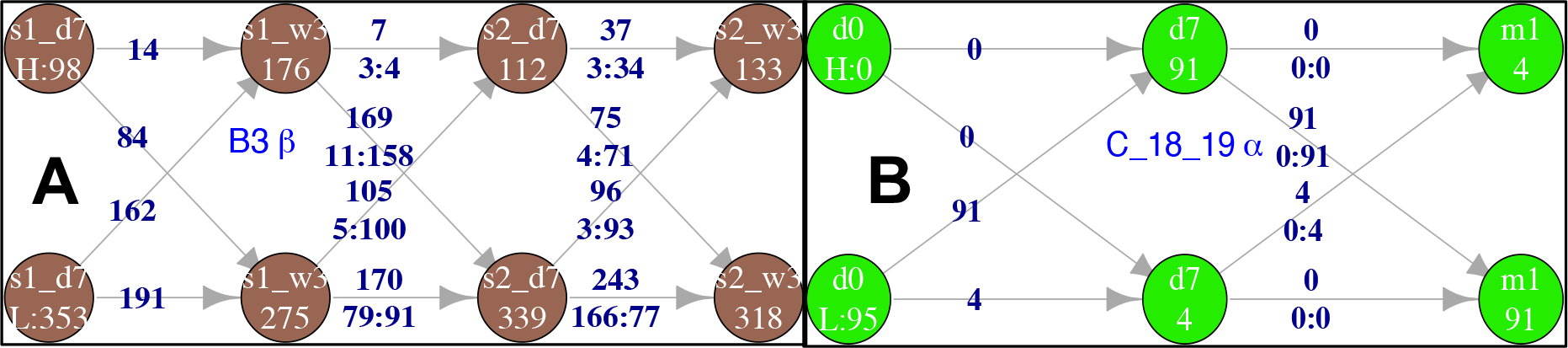
Directed Acyclic Graph for Reactive clones. DAGs (directed acyclic graph) depicting movement of reactive CDR3 clones between the H/L classes across different time points. This data structure is very useful in visualizing the dynamics of clones over time. At each time point a reactive CDR3 can belong to either the H (upper circles) or the L (lower circles) class, and the sum of CDR3s in the two classes is N, the number of reactive clones in an individual. The number on the arrows show how many clones move between circles, the two numbers below that are the number that come from **H:L** circles in the preceding time point (or two steps before the current one). The response has memory, prior states affect the status of clones in subsequent measurements **A)** Covid vaccine data from one donor (B3) for reactive ² clones, using r-fold_m_=10, Δ = 250, τ = 1500. The states are shown for each of the two doses, s1 (dose 1) and s2 (dose2), and the two time points after each dose, d7 (day7), and w3 (week3). **B)** Flu vaccine data from donor C for the 2018-2019 flu season for three timepoints (d0, d7 and m1), using r-fold_m_=5, Δ = 250, τ = 1000. The browser-based tool allows exploration of DAGs.

### The TCR response is an early indicator of the neutralizing antibody response to COVID-19 vaccine

The anti-SARS-CoV-2-spike neutralizing antibody levels in the serum of the COVID-19 vaccine group were measured at 3 time points: week 3 after the first dose (s1w3), week 1 after the second dose (s2d7), and week 3 after the second dose (s2w3). Most individuals (excluding B3 and B4, who were previously infected), exhibited little or no antibody response at week 3 after dose 1 (s1w3), but a robust response at week 1 after dose 2 (s2d7) (**Fig. 4B**), suggesting it takes around 4 weeks for an antibody response to develop, while the T cells start responding already at day 7 after the first dose. Presumably, the antibody values a week after dose 2 (s2d7) reflect the effects of the first dose.

Those cases with a narrow/weak T cell response to the first dose invariably had a weak neutralizing antibody response (B6, B8 and B9) **(Fig. 4)**, and lack of a meaningful T cell response (B9) indicated a very weak antibody response. The correlation between the T cell response (sum of reactive β and β clones) and the antibody response (**Fig. 5**) gives hope that the T cell response is a leading indicator of the immune response. A weak T cell response seems to inevitably presage a weak antibody response (**Table 1, Fig. 5)**, but estimating the antibody response based purely on the T cell response is probably not straightforward because only a subset of the responsive T cells is likely responsible for guiding antibody production.

### Multiple Covid vaccine doses and saturation of the T cell response

The second COVID-19 vaccine dose broadens the immune response in seronegative individuals, inducing new T cell clones, which differ from those induced by the first vaccine dose (s1). However, in individuals recovered from a prior SARS-CoV-2 infection, the T cell response to the second dose (s2) is narrower and weaker than the response to the first vaccine dose. Their antibody levels also decreased at week 3 after the second dose (s2w3) measurement, compared to the measurement at 1 week after dose 2 (s2d7), in contrast to the previously uninfected individuals. The waning T cell response to the second dose in recovered individuals may behave like a third dose would in seronegative individuals.

We assume that there is a fixed number of epitopes from the spike protein that can be presented, dependent on the HLA of the individual. Thus, follow up doses after the first dose elicit responses to shrinking pools of epitopes that have not elicited a response yet, so the response will saturate as this pool of epitopes shrinks.

If we assume that a third dose gets close to exhausting the pool of possible epitopes from the Receptor Binding Domain (RBD), based on our data, the number of possible epitopes varies quite a bit between individuals, and this might possibly explain some of the differences in the TCR response to the vaccine doses.

### Landscape of the T cell response to the Influenza vaccine

For the analysis of the T cell response to the influenza vaccine, we decided to treat the samples from each flu season as a separate “individual” mainly because we had no reason to assume the responses after a year to different flu vaccines would be similar. Indeed, we found no clones responding repeatedly over different flu seasons in any individual, justifying our choice to treat them as separate individuals.

The influenza vaccine samples are annotated with letters (A, B, C, D, and E for the donors) and the flu season (19_20 stands for the flu season bridging the years 2019 and 2020) as well as the timepoint (d7 stands for day 7 and m1 is 1 month after the dose, **Fig. 6**). We note the following features:

a. The α,β response to the flu vaccine is lower, by number of responsive clones, compared to the response to the covid vaccine. Lower thresholds, for r-fold and τ, are needed to identify responsive clones.
b. The α repertoire, is more responsive than the β. This dichotomy between the α and β 40 repertoires is a feature of the response to the Influenza and Covid vaccines in our limited>cohort.
c. Responsive clones at d7 and m1 are distinct (like what was observed in the COVID-19 vaccine group). Responsive T cell clones usually have labels with only one **H**.
d. There is little sign of immunologic memory,
  a. Some individuals respond repeatedly (C), others only responded once (A, B, E), and yet others never at all (D).
  b. Responsive clones are not shared between seasons in an individual

### Symptoms after the vaccine doses and the T-cell/antibody response

Based on our data, we conclude that there is no correlation between the physical symptoms felt immediately after the vaccination dose is administered and the immune response as measured by antibody and responsive T cell clones (**Table 1**), contrary to the popular assumption of a correlation. The symptoms are likely the result of the early innate response, which seems to not be very predictive of the subsequent adaptive response.

## Discussion

### The role of T cells in adaptive immunity against SARS-CoV-2

Mouse models suggest that both humoral and cellular adaptive immunity contribute to viral clearance in primary infections with SARS-COV-2, but the protection against secondary infections or infections after vaccination is largely through antibodies^4^. About 60% of the protection after vaccination against Influenza B in children was determined to be due to antibodies that neutralized Hemagglutination, the rest is potentially due to cellular immunity^13^. These studies were emphasizing the importance of antibodies, but the evidence they cite bolsters the case that the T cell response is a critical component of the protection provided by adaptive immunity. Other studies in mice have demonstrated that poor T cell response underlies severe disease upon SARS-CoV infection^29,30^. T cell dysfunction has also been implicated in poor outcomes in humans infected with SARS-CoV-2^31^. These studies make a strong case for monitoring the T cell clones that respond to vaccines/infections.

### Benefits of T cell monitoring

Longitudinal measurements of T-cell receptor expansions afford a nuanced view of the immune response to the vaccines, in contrast to the measurement of neutralizing antibody response which takes time (weeks) to manifest and provides a single number encompassing the complex response elicited by multiple epitopes.

The benefits of using Tseek on PBMCs to ascertain the response to vaccines are manifold, 1) small non-invasive blood draws are sufficient, 2) can be used within days of a vaccination, 3) enables economic, large-scale analysis of the T cell populations over a time course, in large cohorts, 4) provides highly sensitive and specific results, and 5) has a turnaround time of days.

### The perturbative TCR Repertoire response to vaccination

It is evident from our clustering data that the response to the vaccine is a perturbation of an otherwise relatively constant T cell repertoire; a few hundred clones respond while the rest remain unaffected. An individual’s TCR signature presumably evolves slowly over the years in response to infections.

The clustering together of the two samples post each vaccine dose suggests weeks-long persistence of the responsive clones after the initial perturbation. This perturbative response is probably the key indicator of a healthy response, too strong a response might be a sign of dysfunction and could be monitored for remedial action.

### Breadth of the T cell response

The “breadth” of the response to vaccines/infections, how many viral epitopes are targeted in the response, is probably a key determinant of how well the adaptive response protects against future infections by the virus or its variants. This is a subjective measure, a comparison between the immune responses to the Influenza and Measles viruses provides some context.

The response to influenza infections is polyclonal, but is defined as narrow because the virus can escape immune surveillance in an individual with a single mutation^32^. Individuals differ in the dominant epitopes, leading to private viral mutations for escape, unlike in ferrets, which get broad immunity after an infection by the Influenza virus. Prior infections by Influenza can lead to pre-existing immunity mounting responses to conserved viral epitopes from early strains, the “antigenic sin antibodies”, which are often non-neutralizing epitopes^33^. These make frequent vaccinations against influenza a necessity.

In contrast, the same Measles virus vaccine has been used for decades and the virus has not evolved mutations that can evade the vaccine-induced immunity, because the response includes multiple co-dominant B cell epitopes from the surface glycoproteins, making escape from immune surveillance difficult, by requiring coordinated changes in multiple sites (at least 3), which is extremely unlikely^34^.

Broad T cell responses targeting multiple dominant epitopes from viral proteins are an effective insurance against viral mutations escaping immune surveillance. Our TCR repertoire data from the flu and covid vaccines suggests the T cell response is broad in the case of the Covid-19 mRNA vaccines, in contrast to the T cell response to the influenza vaccines.

### Response to influenza vaccine

The number of T cell clones responsive to the influenza vaccines, even in responsive individuals, is lower than in the case of the COVID-19 vaccine. This could arise from the difference in vaccine platform with the influenza vaccine using inactivated influenza virus while the COVID-19 vaccine is using the mRNA of SARS_CoV-2 spike protein.

The **β** chain repertoires have quite a narrow response to the influenza vaccine, while the **α** chain repertoires are broader. The TCR response to the influenza vaccine suggests the importance of monitoring both the TCR **α** and **β**chain repertoires. It is not clear what implications an imbalance in the response of the two chains has for immunity; it would be interesting to study the ferret’s T cell response to the influenza virus, since ferrets seem to get immunity to influenza similar to the human response to measles.

Some individuals in our study respond to vaccinations over consecutive flu seasons, while others never respond, suggesting that the differences in response are individual, overriding any effects of immunological memory,

### Response to Covid vaccine

For the covid vaccine responses, multiple reactive clones arise at different times over a period of a month or longer. Studying this in detail requires frequent measurements which are impossible to conduct in humans, due to obvious ethical reasons.

One possible explanation is that these diverse clones target the same epitopes. A recent study on Yellow fever virus (YFV) vaccination^35^ suggests that of the multiple clones that have binding affinity to an epitope, only a few get selected for expansion, probably in a stochastic manner. This is unlikely to be the case here, as the immune response at any given time will eliminate cells presenting epitopes they already recognize, potentially preventing multiple clones against the same epitopes arising in a single response. Therefore, we believe that new responsive clones likely target new epitopes. All possible viral epitopes (compatible with the individual’s HLA) continue to be presented over time and cells presenting epitopes already recognized by existing, expanded T cell clones probably get eliminated, discouraging redundancy. This extended response diversifies the response, and ensures any lingering infection gets a robust response.

Another possible explanation for the diversity of clones is the two wave theory^36^, which postulates several classes of T cells with complicated dynamics. But the simplest explanation is that there is a constant turnover which more frequent sampling would reveal, and the appearance of seemingly distinct populations is a quirk of the timing of our measurements.

The mostly new clones responding after the second dose suggest that recently stimulated T cells in response to the first dose probably cannot clonally expand again. Such a refractory period for T cell stimulation might prevent overstimulation and broaden the response over many T cells and epitopes.

Even though the receptor-binding domain (RBD) is immunodominant^37^ and 90% of the neutralizing activity present in sera post SARS-CoV-2 infection targets the RBD, the diversity of reactive clones suggests a large number of epitopes contribute to this immune protection^38^, which cannot be discerned from the measurement of the activity of neutralizing antibodies.

### Vaccine versus infection

Differences in the adaptive immune outcomes of vaccination versus infection provides a window into the underlying mechanisms. In the case of influenza, vaccination induced reactions to neutralizing epitopes^33^; infections on the other hand induced non-neutralizing responses to conserved viral epitopes from early strains (the antigenic sin antibodies). This might be specific to influenza in humans in contrast to ferrets.

In humans, SARS-CoV-2 infections have been shown to elicit higher levels (compared to mRNA vaccines) of original antigenic sin antibodies that bind more strongly to other seasonal coronaviruses compared to SARS-CoV-2^39^. This would suggest that the vaccines are “superior” to natural infections. Another line of reasoning suggests vaccination provides better protection than infection by SARS-CoV-2, because antibodies generated following a SARS-CoV-2 infection also target non-neutralizing viral targets, while the mRNA vaccines elicit a focused, neutralizing response^40^. However, the finding that the spike RBD is immunodominant (90% of the neutralizing activity present in sera post SARS-CoV-2 infection targets the RBD) suggests that the reality is more complicated than the simple conclusion that immunity conferred by infection is “inferior” to that conferred by vaccination^38^.

Additionally, the order of infections by different SARS-CoV-2 strains might be important in determining the protection against variants. One study showed that parental alpha strains induce antibodies that cross-react with other strains (B.1.1.7, B.1.351) while an initial infection by B.1.1.7 resulted in antibodies that were significantly worse in recognizing and neutralizing the parental strain^41^.This suggests that vaccines targeting specific sub-types need not be better.

Based on our limited data, infection by SARS-CoV-2 is equivalent to a mRNA vaccine dose, and both likely provide equivalent protection against variants, due to a broad response.

### Cross-reactive T cells and indicators of resilience in the repertoire

The variable outcomes of infection by SARS-CoV-2 led to suggestions that this variability is due to the existence of protective, cross-reactive CD4 T cells in some responders. Several studies have ruled this out, finding no differences in neither the TCR repertoire^42^ nor the antibody responses^43^ between patients with mild and severe cases of COVID-19 in patients. The YFV vaccine study found pre-existing cross-reactive T cell clones against Yellow Fever, but they were not the ones reacting to the YFV vaccine^35^, suggesting cross-reactive clones, even if they exist, might not offer readymade protection against infections. This suggests prediction of the response based on T cell clones already present might not be possible.

### Effect of prior exposure to SARS-CoV-2 on the vaccine immune response

A study^44^ showed that individuals recovered from SARS-CoV-2 generated higher antibody titers after the first dose than did those without prior infection by using RBD ELISA (for the antibody response), and interferon-γ release assays and intracellular flow cytometry (for B and T cell response). The antibody titers following the first dose in recovered individuals was on par with those seen in the second dose in seronegative individuals. The T cell response was also determined to be stronger in the convalescent, but this was measured indirectly, through ELISpot. Using mouse models, another study suggested that both humoral and cellular adaptive immunity contribute to viral clearance in primary infections, but the protection against secondary infections or infections after vaccination is largely through antibodies^4^.

We find that the T cell response to the first dose is similar in the recovered, and the pre-existing antibodies also get a boost, but, after the second dose, the antibody levels seem to go down, along with the T cell response. The benefits from additional doses for these individuals will probably be limited and prior exposure is likely equivalent to the first vaccination dose for the uninfected.

### Efficacy in population

The reported efficacies of the mRNA vaccines have been in the 85-90% range. Our limited data suggests that 1 in 9 might have a limited response to the vaccine which seems consistent with the epidemiological data.

### Booster doses

Most responsive clones after the second dose are new, not seen before in the responses to the first dose. The distinct response to the second dose broadens the response, making it more resilient against point mutations.

A third dose might not always end up boosting antibody levels, as suggested by our limited data, where recovered individuals exhibited a decrease in antibody levels 3 weeks after the second dose. But a third dose might be essential for individuals who mount a weak response after two doses (like B9 in our cohort).

### Spacing between the doses

Our data suggests that the response to the first dose is ongoing, with new T cell clones still being generated, at the time of administration of the second dose in the current dosing schedules (3/4-weeks for Pfizer/Moderna vaccines). Increasing the spacing between the first and second doses might allow the response to the first dose to play out, before further immune stimulation and give better protection, several studies now suggest this ^45,46^. Data from Canada and Great Britain also seem to suggest that increasing the spacing might give better protection^47^. An alternate possibility is that administering the second dose close to the first one might broaden the T cell response by inducing new clones that don’t overlap with the first response, only a controlled large-scale study can provide a definitive answer to this question.

### Waning response

It will be difficult to see the responsive T cells after 6 months, and the levels of antibodies might wane, but the memory B and T cells probably persist and help maintain responsiveness. Supporting this persistence of memory B/T cells, a recent study showed vaccines continued to protect against severe disease at close to 100% levels, despite the reduction in circulating antibodies over time^48^.

### Caveats

The fundamental assumption in our analysis is that the abundance of vaccine reactive TCR **α** and **β**clones corresponds to binding efficiency to their cognate viral epitope. Implicit in this assumption is that there is an approximately one-to-one mapping between TCR CDR3 and epitopes. Clustering CDR3 by homology (Methods) did not affect our trees, nor did it significantly change the number of reactive clones but clustering in this manner might also not be sufficient in any case, since it might not identify all clones with common epitope targets, as there are cases where non-homologous clones that share targets, and homologous clones that have completely different targets.

## Conclusions

Our data suggests the TCR repertoire is a useful and meaningful biomarker for the study of immune reaction to the vaccine. The TCR repertoire adds an additional dimension, providing a detailed view of the effect of vaccinations on the immune system that has the advantage of being non-invasive, inexpensive, and easy to implement. A study based on small cohorts does not allow inference of broad truths, but certain observations, such as the fact that the symptoms just after the vaccination dose are not indicative of the adaptive immune response are likely to be broadly true. This study is a start, additional large-scale studies are needed to fully realize the potential of these measurements. The accompanying website, http://katahdin.girihlet.com/shiny/tvax/, allows full exploration of the data.

## Methods

### Experimental

#### Tseek

The T cell receptors (TCR) are heterodimers of an α and a β chain, which are trans-membrane polypeptide chains linked to each other by covalent disulfide bonds. Hyper-variable complementarity determining regions (CDRs), located on the α and β chains of V regions of the TCR determine the binding specificity of the receptors to peptide-MHC complexes. Of the three CDRs in the α and β chains, CDR3 is formed by somatic rearrangements of V and J (joining) gene segments in the α chain or V, D (diversity) and J segments in the β chain, during the maturation of the T lymphocytes. The CDR3 contains additions and deletions of nucleotides that are not coded in the genome. The organization of the a and b loci in the mouse and human genomes are depicted in Fig. S1.

Briefly, total RNA from PBMC’s from 2 ml of blood was extracted using AllPrep DNA/RNA Micro Kit from Agilent Technologies (Cat. No.: 80284). The quality and quantity were checked using the RNA Bioanalyzer Nano (Agilent Biotechnologies). Messenger RNA (mRNA) from 500ngs of total RNA was isolated and fragmented, and reverse transcribed using random primers. Adapters with 8bp molecular indices were ligated to all the blunt ended fragments. Nested PCR was then performed using a TCR constant primer and an adapter primer. Libraries were quantified again using a bioanalyzer and sequenced on Illumina Next seq 500 (150bp PE). CDR3 peptide sequences were identified, and the frequencies were tabulated in all the samples.

#### Spike antibody measurements in serum

The kit used here uses a protein-based surrogate neutralization ELISA (snELISA) approach, which measures the ability of antibodies to prevent the association of soluble biotinylated ACE2 to immobilized RBD: A higher signal (snELISA integrated score) in this assay indicates low neutralization^49^.

Vaccine-elicited anti-SARS-CoV-2 antibody responses were quantified using the GenScript SARS-CoV-2 Neutralization sVNT cPASS TM Kit (GenScript Catalog #L00847-5), which effectively measures the ability of patient plasma to block the interaction between the receptor binding domain (RBD) of the SARS-CoV-2 spike protein and its receptor angiotensin converting enzyme-2 (ACE2), according to manufacturer’s instructions. Briefly, blood was collected by venipuncture into Vacutainer EDTA tubes (Becton Dickinson Catalog # 368589), and centrifuged at 1000xg for 10 minutes at 4°C. The upper, clarified plasma fraction was transferred to a new tube and spun again at 1000 x g for 10 minutes at 4°c to remove residual cellular material and debris. The plasma was aliquoted and snap frozen at minus 80°C. For testing, the plasma was thawed on ice, and a dilution series was generated by diluting in Sample Dilution Buffer. Each dilution was incubated with recombinant horse radish peroxidase (HRP)-conjugated RCB for 30 minutes at 37°C to allow antibody/RCB binding, transferred to human ACE2 coated assay microtiter plates, and incubated for 15 minutes at 37°C for 15 minutes. Plates were washed extensively (4x), and 100ul of TMB solution was added to each well, and plates incubated in the dark for 15 minutes at 25°C. Stop Solution was added to each plate, and the absorbance of each sample was immediately read in a microtiter plate reader at 450nm. Sample absorbances were compared to the neutralization curve of a commercially available anti-SARS-CoV-2 RCB monoclonal antibody (GenScript, Catalog # A02051) of known concentration to extrapolate antibody titers, according to manufacturer’s protocol.

### Analytical

#### Processing Tseek data

The data from Tseek is nucleotide sequences that start in the V segment, span the CDR3 region and cover the whole J segment in β and β chains. The sequencing reads are merged for paired-end data and mapped to the V and J segments (based on annotations from Gencode^50^). Reads that are annotated with V and J identities are translated to amino acid sequences (all three frames), frames with stop codons are filtered out. Based on the V and J annotations, the location of the CDR3 region is identified, and the precise boundaries are determined using motifs for the boundaries derived from annotated sets of CDR3, which is based on a reference set in the IMGT database^51^. Separate tables are created for the V and J segments, the V-J pairs, and CDR3 counts, across all samples in the study. The rest of the analysis is based on these master tables.

#### Analysis of CDR3 data

The methods described here are also implemented in tools on the companion website to this study, (http://katahdin.girihlet.com/shiny/tvax/) which allows users to try different settings and values to explore the data and generate the figures and tables shown in the paper.

#### Comparisons of two distributions

Entropy (H) for a vector with N probability members is given by - Σ_i_p_i_ log(p_i_) where, Σ _i_p_i_ = 1. The maximum entropy possible is log(N), which is used to normalize the entropy.

Grouping/clustering samples requires sample comparisons and measuring a “distance” between them. Two metrics that are proper distances are the Weighted Jaccard (Jw) and the Shannon-Jensen (JS) distance/divergence^52^, which can be used to build trees. The JS divergence between two distributions and, is defined as

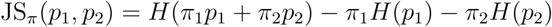

where, the π are the relative weights of the two distributions, satisfying π_1_,π_1_ > 0, π_1_ + π_2_ =1

#### Definition of responsive clones

T cells with TCRs that recognize MHC-presented peptides from antigens receive signals upon binding to expand clonally. We identify clonal expansion by calculating the fold change in numbers over the baseline keeping in mind that we are measuring the changes in PBMCs, which is remote from the site of expansion.

To avoid division by zero and high fold-changes for low-expressed clones, we calculate fold-change using the formula, r-fold = (max + Δ) / (min + Δ), where max/min are the maximum/minimum frequencies of the clone in all the samples and the Δ ensures low-expressed clones don’t dominate the list of reactive clones, e.g. using Δ = 100 for a clone whose frequency changes from 1 to 100 (100-fold change) results in r-fold ∼ 2.

We define an expression threshold, τ, values above this are labelled high (H) and below this are low (L). Reactive clones are those whose maximum value is above τ and whose r-fold > r-fold_m_. This definition also implicitly depends on Δ.

We did not always manage to get samples from each donor before the first vaccine dose. Using the criteria listed above identifies reactive clones in the case of donors for whom we have the pre-vaccine data, and by interpolation, we believe it also holds for the samples with missing pre-vaccination data. For the covid samples, we chose Δ_C_ = 250, r-fold_Cm_ = 10, and τ_C_ = 1500 while for the flu samples, recognizing a weaker response, we picked Δ_F_ = 250, r-fold_Fm_ = 5, and τ_F_ = 1000.

#### Clustering CDR3

We used clusTCR^53^ to cluster TCRs based on CDR3, ignoring the V, J labels, as well as HLA information. clusTCR uses a two-step clustering approach, using Faiss Clustering Library for speed and Markov Clustering Algorithm (MCL) for accuracy. The clusTCR publication has compared the results to outputs from GLIPH2^54^, iSMART^55^, and tcrdist3^56^, showing it has comparable clustering quality with improvements in speed and scalability. Clustering had negligible impact on our results.

### Ethics approval and consent to participate

#### Covid vaccine cohort

The Institutional Review Board (IRB) of the Mount Sinai School of Medicine Icahn reviewed and approved the protocols, informed consent, and other study documents (HS #: STUDY-21-00050) for collecting blood samples through venipuncture from consenting healthy, non-pregnant adults who have received or will receive vaccination against covid.

#### Flu vaccine cohort

The anonymized flu-vaccine samples were obtained from the flu-vaccine biorepository at Mount Sinai, headed by Dr. Viviana Simon.

## Abbreviations

TCR: T cell receptor
CDR3: Complementarity-determining region 3
HLA: Human Leukocyte Antigen
PBMC: Peripheral blood mononuclear cell
VDJ: Variable-Diversity-Junction
YFV: Yellow fever Virus
mRNA: messenger Ribonucleic acid
RBD: Receptor-Binding domain
ELISA/ELISpot: Enzyme-linked immunosorbent assay/spot

## Acknowledgements

Stu Aaronson, Glaucia Furtado and April Davis gave many critical comments. We thank the anonymous donors for their dedication to the project and their diligence in strictly following the schedule for blood draws. RS acknowledges the antibody engineering/phage display course at CSHL for insights into antibodies.

