## supplementary material for "The T cell receptor repertoire reflects the dynamics of the immune response to vaccination" (from another version of this preprint: version 1, 2021-12-12)

By Mohammed et. al

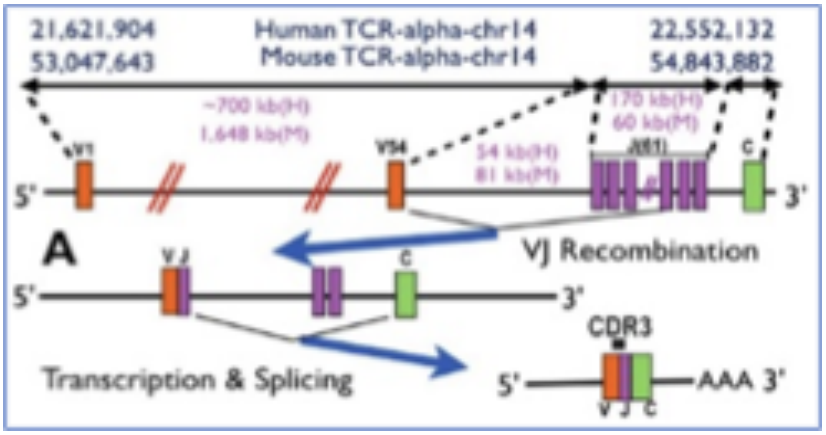


**Figure S1. TCR loci in mouse/human genomes. A)** α locus. Multiple choices for V (orange, ~100 segments, ~400 nt long), and J (violet, ~50 segments, ~65 nt long), and only one C (green, 4 exons, ~1000nt long). A V and a J combine in the genome (DNA) to create the CDR3, while the J-C junction is formed by mRNA splicing. **B)** β locus. Multiple choices for V (orange, ~34 segments, ~400 nt long), two D's (blue, ~15nt long) and two groups of J and C. The first J, C combination consists of J1 (violet, 6 segments, ~50 nt long), C1 (green, 4 exons, ~1000nt long), and the second consists of J2 (violet, 7 segments, ~50 nt long) and C2 (green, 4 exons, ~1000nt long). V, D and J recombine in the genome (DNA) to create the CDR3; mRNA splicing creates the J-C junction. The final transcript is one of two combinations: V/D1/J1/C1 or V/D2/J2/C2.

### The Covid spike protein

The Covid vaccine targets the SARS-CoV-2 spike protein (1273 aa long) consisting of an N-terminal signal peptide (1–13), the S1 subunit (14–685) responsible for receptor binding, and the S2 subunit (686–1273) responsible for membrane fusion. The S1 subunit has an N-terminal domain (14–305) and a receptor-binding domain (RBD, 319–541); the S2 subunit has a fusion peptide (FP) (788–806), heptapeptide repeat sequence 1 (HR1) (912–984), HR2 (1163–1213 residues), TM domain (1213–1237), and a cytoplasm domain (1237–1273). The RBD is a dominant epitope, at least neutralizing antibodies often target this[37] but there might be other epitopes that also induce a response that provide ample protection against future infections.

### DAG model

At each time point, a responsive segment can be labelled as either low (**L**) or high (**H**). **H** and **L** form a binary code, for ***n*** time samples there are $2^n$ possible label strings (**LLLH**, **HLLL** etc. for 4 samples). The table of *Label counts* (Tables 1 and 2) lists counts for all possible label strings.

The DAG figure has two circles for each sample, a high circle that contains all the H segments and the low circle that contains all the L segments. Arrows connect each circle to the two circles in the following timepoint. A sample can only move into one of two circles (H or L) at a subsequent time point. The arrow shows numbers, below each number are two numbers that show the *history*. The first number in history is the number that came from a high circle in the previous time point and the second number is the part that came from the low circle in the previous time point. This shows if there is any *memory* in the process, where a prior state affects the subsequent state. This is another way of asking if the process is Markovian (no memory of previous state) or not. In this case the process does exhibit memory, a clone that is H earlier, mostly transitions to L at subsequent timepoints.

### Representation of the response.

Matrices and transition probabilities can be used to model the DAG. The **N** responsive CDR3 are partitioned into two sets, **H** (high) and **L** (low) depending on frequencies being above or below a user-specified threshold. A matrix $M_{12}$ can be used to study the transitions from samples at timepoint 1 to samples at timepoint 2, with the matrices defined as follows,

$$M_{12} = \begin{bmatrix} \mathrm{p}_{HH}^1 & \mathrm{p}_{LH}^1 \\ \mathrm{p}_{HL}^1 & \mathrm{p}_{LL}^1 \end{bmatrix} = \begin{bmatrix} \mathrm{p}_{HH}^1 & 1-\mathrm{p}_{LL}^1 \\ 1-\mathrm{p}_{HH}^1 & \mathrm{p}_{LL}^1 \end{bmatrix}$$

$$\begin{bmatrix} p^1_{HH} & 1-p^1_{LL} \\ 1-p^1_{HH} & p^1_{LL} \end{bmatrix} \begin{bmatrix} H_1 \\ L_1 \end{bmatrix} = \begin{bmatrix} H_2 \\ L_2 \end{bmatrix} = \begin{bmatrix} p^1_{HH} * H_1 + (1-p^1_{LL}) * L_1 \\ (1-p^1_{HH}) * H_1 + p^1_{LL} * L_1 \end{bmatrix}$$

where, $p^1_{HL}$ stands for the probability a clone in the state “H” at the first time point (1) transitions to an “L” at the subsequent timepoint.

$M_{13} = M_{23} * M_{12}$ is satisfied only if there is no dependence on history, if the process is truly Markovian. Despite the non-Markovian nature of the history, we can use the values for the matrix inferred from our data to estimate how many T cells react to the vaccine on average, at each measurement.

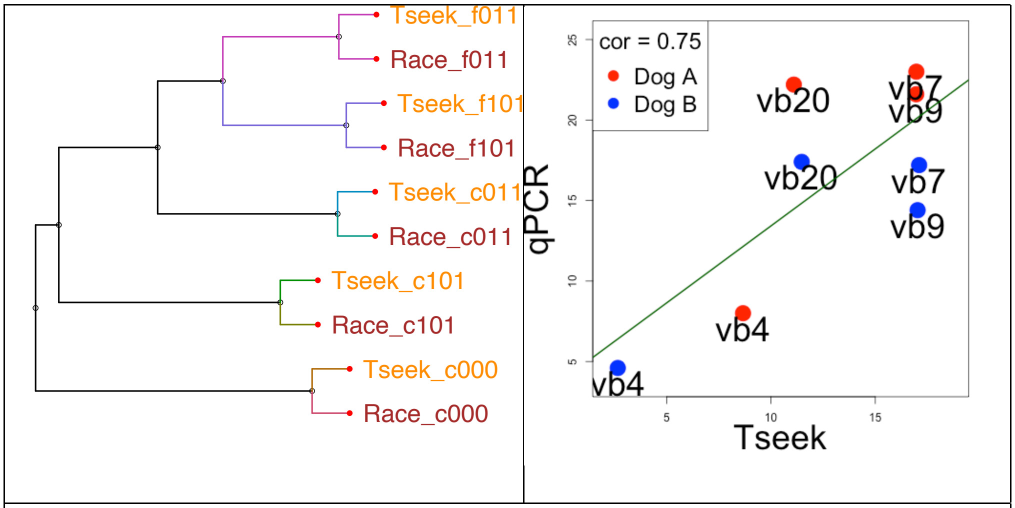


**Figure S2 A) Comparison of Tseek and RACE-PCR.** TCR profiles were generated from the same set of samples using Tseek and RACE-PCR. A dendrogram constructed using Jensen-Shannon divergence for $\beta$ CDR3 is shown in the figure, $\alpha$ has a similar tree. PBMC samples were either untreated or treated with flu M1 protein or flu M1 conjugated with CD40 that signal dendritic cells to process the proteins. The “c” stands for stimulation with just the Flu M1 protein, while the “f” stands for samples stimulated with the Flu M1 protein conjugated with CD40. 011 and 101 are the concentrations (nM) of the M1 protein, c000 is the unstimulated sample. The clustering of the data suggests RACE-PCR (Race) and Tseek generate similar profiles, but Tseek has higher sensitivity (can start with smaller amounts of material) and specificity (150 nt reads are sufficient to identify V, J and CDR3 and most reads are from the TCR). Other methods of profiling TCR repertoire use multiple V primers for PCR, resulting in various biases needing complex corrections. **B) qPCR versus TCR.** In a dog data set, primers for qPCR were designed for four V segments with different expression levels. RNA from PBMCs of two dogs (Dog A, red and Dog B, blue) were processed using T seek to determine the expression of the V segments, in parallel qPCR was used to determine levels of the V segments in the same samples. The plot shows qPCR results (y axis, cycles compared to the reference) against abundances from Tseek (x axis, log2 scale). The abundances vary over 2 orders of magnitude and show good concordance.

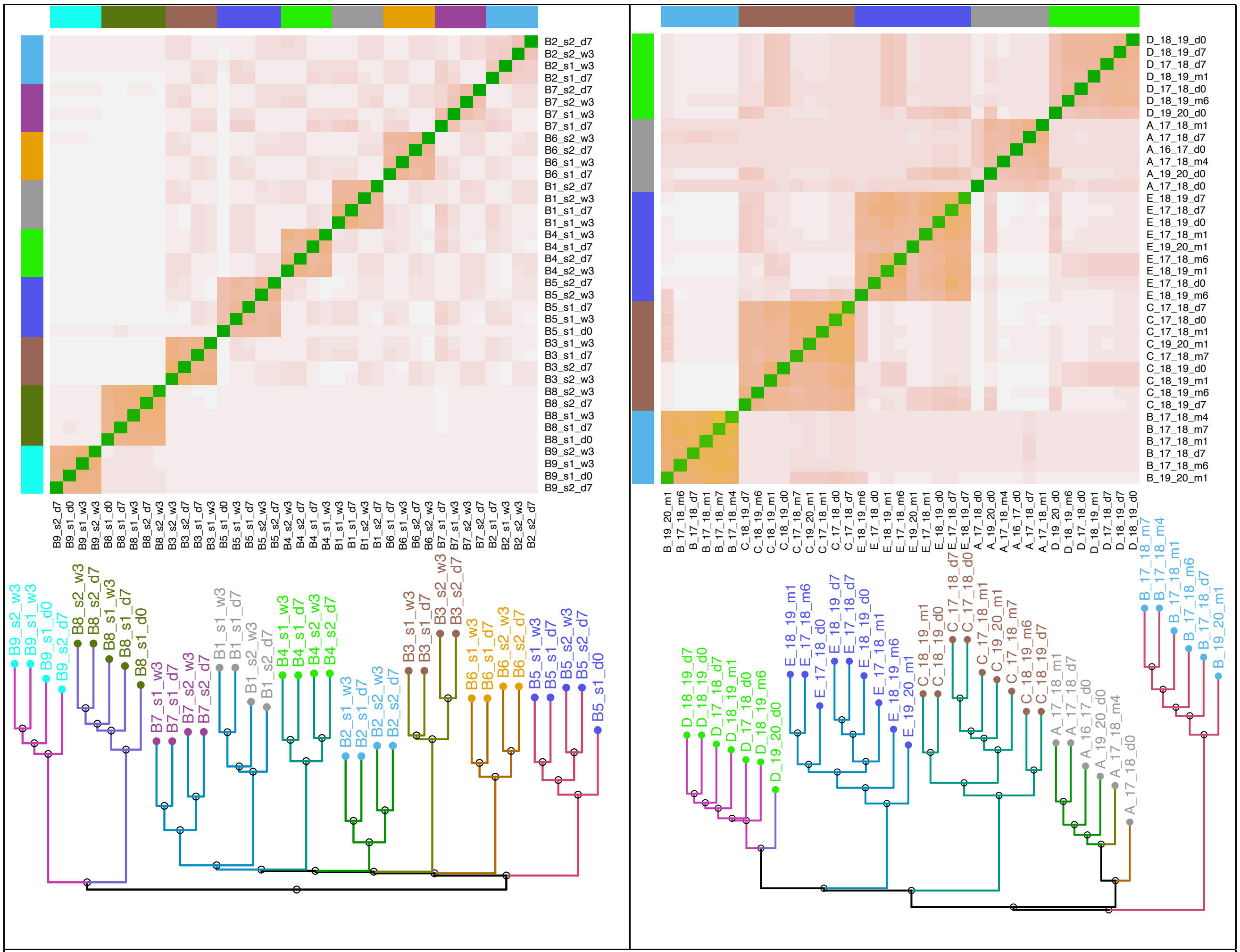


**Figure S3 Clustering of Flu/Covid vaccine samples based on CDR3 β.** Distances between samples were defined for the CDR3 data using the Jensen-Shannon metric (an information-theoretic measure, Supplementary material). The heatmaps show the distance between pairs of samples, while the dendrograms show clustering of samples by individual. **A) Covid CDR3 β.** The first dose is s1, the second dose is s2, d0 is the pre-vaccine sample, d7 is 7 days post vaccination, w3 is 3 weeks post vaccination. For each individual, the two samples after the second vaccine dose (s2_d7 and s2_w3), cluster separately from the two samples after the first vaccine dose (s1_d7 and s1_w3) suggesting distinct sets of T cells react to the two doses. The second dose broadens viral epitopes targeted, likely improving the immunity provided by the vaccination. (**Fig. 3A** shows the corresponding $\alpha$ tree) **B) Flu β CDR3** 19_20 refers to the 2019-2020 flu season, d0 is the pre-vaccine sample, d7 is 7 days post vaccination, m1 is 1 month post vaccination. The clustering by vaccine dose does not occur in some individuals (B, D), and the m4 and m6 samples seems to have lost "memory" of the dose, the T cell response does not persist beyond 2-3 months. Even though there are responsive clones for each flu season (Table 2A, B), the relative lack of clustering of the tree by vaccine dose, in contrast to the covid vaccine data suggests that the response is "weaker" and has individual and seasonal differences. **Fig. 3B** shows data for the $\alpha$.
